# Tricarboxylic acid (TCA) cycle, sphingolipid, and phosphatidylcholine metabolism are dysregulated in *T. gondii* infection-induced cachexia

**DOI:** 10.1101/2022.12.20.521281

**Authors:** Tzu-Yu Feng, Stephanie J. Melchor, Haider Ghumman, Mark Kester, Todd E. Fox, Sarah E. Ewald

## Abstract

Cachexia is a life-threatening disease characterized by chronic, inflammatory muscle wasting and systemic metabolic impairment. Despite its high prevalence, there are no efficacious therapies for cachexia. Mice chronically infected with the protozoan parasite *Toxoplasma gondii* represent a novel animal model recapitulating the chronic kinetics of cachexia. To understand how perturbations to metabolic tissue homeostasis influence circulating metabolite availability we used mass spectrometry analysis. Despite the significant reduction in circulating triacylglycerides, non-esterified fatty acids, and glycerol, sphingolipid long-chain bases and a subset of phosphatidylcholines (PCs) were significantly increased in the sera of mice with *T. gondii* infection-induced cachexia. In addition, the TCA cycle intermediates α-ketoglutarate, 2-hydroxyglutarate, succinate, fumarate, and malate were highly depleted in cachectic mouse sera. Sphingolipids and their de novo synthesis precursors PCs are the major components of the mitochondrial membrane and regulate mitochondrial function consistent with a causal relationship in the energy imbalance driving *T. gondii*-induced chronic cachexia.

## Introduction

Cachexia is a metabolic disease defined by loss of 5% or more lean body mass in under 6 months with or without adipose tissue loss, anorexia, weakness, fatigue, and systemic elevation of inflammatory cytokines or tissue damage markers^1^. Cachexia is a common comorbidity of a wide range of chronic diseases including cancers, chronic heart failure, chronic obstructive pulmonary disease^2^, chronic liver and kidney diseases^3^, rheumatoid arthritis^4^, and chronic infections^5,6^ In 2014 cachexia was estimated to affect 9 million people in the United States, Europe, and Japan^7^ Patients with cachexia have a poor quality of life and worse outcomes compared to non-cachectic patients. For example, Orell-Kotikangas *et al*. reported that cachexia is associated with shorter disease-free survival in head and neck cancer (13 months vs. 66 months)^8^. In addition, cachectic patients have prolonged hospitalization leading to higher health care costs^9^. Although appetite stimulants and anti-inflammatory agents have been explored as cachexia therapies, these treatments have not been broadly efficacious for improving quality of life or reducing mortality^10^.

The pathology of cachexia is thought to be driven by a hypercatabolic energy imbalance. Increased muscle proteolysis and decreased protein synthesis in skeletal muscle and cardiac muscle are the hallmarks of the disease^11^. Unlike muscle wasting, the morphological and functional changes in adipose tissues can be heterogeneous during the development and progression of cachexia. It has been proposed that adipose atrophy in cachectic patients is due to increased lipolysis and fat browning^11^. Paradoxically, increased adipocyte lipogenesis has also been observed in cachectic human patients and murine models^12–15^. In addition to dysregulated adipose tissue and muscle loss, impaired liver functions are observed in cachectic patients and animals^16,17^ Aberrant liver functions can provoke imbalanced energy metabolism in cachectic animals^18^. These, sometimes conflicting reports, highlight the need for better animal models that will lead to a mechanistic understanding of the metabolic dysregulation in cachexia and facilitate the development of novel therapeutic strategies^19^.

Current animal models fail to recapitulate the chronic nature of clinical cachexia. For example, surgical and tumor models of cachexia have a brief window between the onset of weight loss (anorexia cachexia) and the time point when animals succumb to the initiating insult^20,21^. To more accurately recapitulate the immune and metabolic disruptions of chronic cachexia there has recently been renewed interest in establishing infectious disease models of cachexia. Cachexia is frequently observed in protozoan parasite infections including Malaria (*Plasmodium sp.*)^22^, Leishmaniasis^23^, African Trypanosomiasis (*Trypanosoma bruceĩ*)^24^ and Chagas Disease (*Trypanosoma cruzi*)^25^. Work from our lab^15,16,26^, and others^27–29^, has recently established Toxoplasmosis as a robust model to study the regulation of acute and chronic cachexia in mice. *Toxoplasma gondii* is an obligate intracellular parasite that naturally infects mice and humans, establishing life-long infection in neurons, skeletal and cardiac muscle. We have previously shown that intraperitoneal or oral infection with *T. gondii* can result in chronic cachexia characterized by acute anorexia and adipose tissue loss followed by sustained skeletal muscle wasting, elevated serum inflammatory cytokines, and persistent weight loss^15,16,26^, rendering a tool to study metabolic changes during chronic cachexia.

This study aimed to understand perturbations to circulating metabolites in mice with *T. gondii*-induced chronic cachexia. *T. gondii*-induced chronic cachexia was associated with a significant reduction in the weights of muscle and liver but not subcutaneous and visceral adipose tissues, consistent with previous observations that adipose tissue wasting is not necessary for cachexia. Using untargeted mass spectrometry-based metabolomics, we determined that several sphingolipids and glycerophospholipids are enriched in the sera from cachectic animals as compared to non-cachectic controls. Sphingolipids are one of the major components of the membranes of cells and organelles, including mitochondria. The dysregulation of sphingolipid metabolism significantly impairs mitochondrial functions in yeasts^30^ and mammalian cells^31^. Sphingolipid-targeted mass-spectrometry analysis showed that cachectic mice have increased C16 sphingomyelin (SM) species, which have been linked to cancer-induced cachexia^32^ and liver diseases^33,34^ In addition, the levels of multiple intermediates of trichloroacetic acid (TCA) cycle, including α-ketoglutarate, 2-hydroxyglutarate, succinate, fumarate, and malate, were reduced in *T. gondii*-induced chronic cachexia compared to uninfected, non-cachectic mice. Collectively, these data suggest that liver atrophy-associated sphingolipid metabolism dysregulation may induce aberrant mitochondrial functions resulting in defective TCA cycle activity in *T. gondii*-induced chronic cachexia.

## Results

### *T. gondii* infection-induced chronic cachexia is associated with decreased serum lipids

Consistent with previous reports, mice intraperitoneally infected with 10 *T. gondii* cysts lost, nearly 15% of their body weight in the first 2 weeks of infection (Fig. 1A)^15,16,26^. Weight loss was sustained for 9 weeks, when the experiment was terminated. Parasite burden at 9 weeks postinfection was confirmed by quantitative polymerase chain reaction (qPCR) for *T. gondii RE* in brain genomic DNA. (Fig. 1B). At 9 weeks post-infection skeletal muscle mass, measured by quadriceps weight, (Fig. 1C) and liver mass (Fig 1D) was significantly lower in infected mice relative to uninfected controls. Inguinal subcutaneous white adipose tissue (scWAT) and epigonadal visceral white adipose tissue (vWAT) were not significantly different between infected mice compared to age-matched controls (Fig. 1C). This result is consistent with previous studies showing adipose tissue loss during the acute, anorexic phase of *T. gondii* infection, (5-14 days post-infection), however, as mice regain eating adipose mass rebounds^16,26^. These studies also found that chronic cachexia was not associated with increased non-shivering thermogenesis, fat browning, or lipolysis^15^. However, the levels of circulating lipid species in cachectic mice were not investigated.

**Figure 1.**
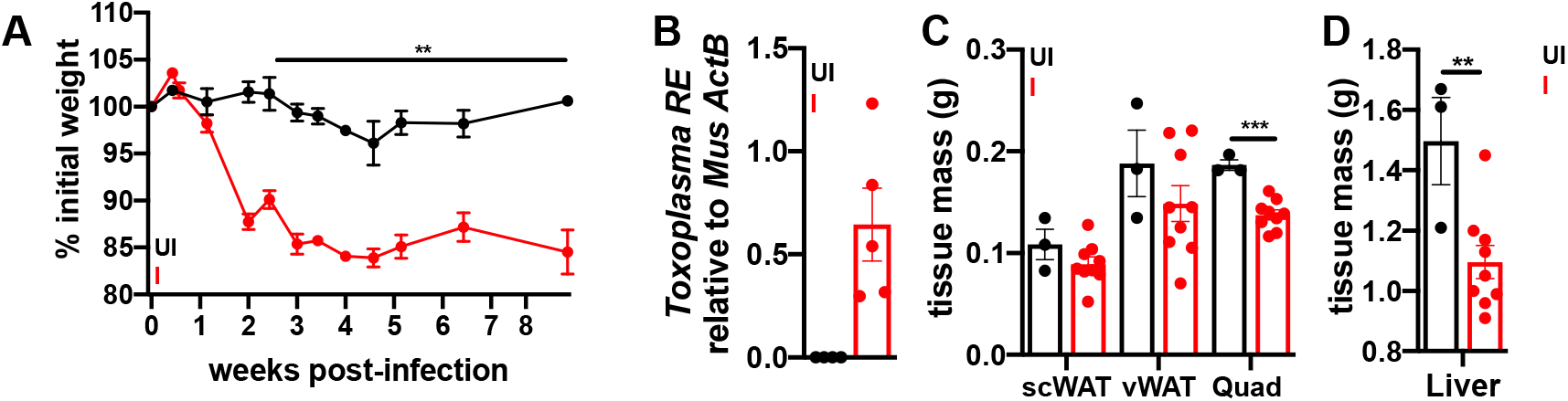
*T. gondii* infection causes chronic cachexia in mice. 10-14 week old C57BL/6J mice were intraperitoneally infected with 10 Me49gLuc *T. gondii* cysts of the Me49 strain or mock injected with PBS. A, Weight curves represent change from initial weight in uninfected (UI) and infected (I) mice. Asterix represents time points where UI vs. I weights are significantly different. B, Qpcr for *T. gondii* RE genomic DNA normalized to mouse beta-actin in brains from chronically infected mice 9 weeks post-infection (wpi) or uninfected littermate controls. C-D, At 9 wpi inguinal subcutaneous white adipose tissue (scWAT), epigonadal visceral white adipose tissue (Vwat), quadricep muscles (Quad) ©, and liver (D) were dissected and weighed. N = 3-9 mice per group, representative of 3 independent experiments. Error bars represent the mean+/-SEM \**p*<0.05, \*\**p*<0.01, \*\*\**p*<0.001 by unpaired Student’s t-test.

During lipolysis, triacylglycerides (TAGs) are broken into non-esterified fatty acids (NEFA) and glycerol which are released at high levels into circulation^35^. While cachectic mice had low serum TAGs (Fig. 2A), NEFA (Fig. 2B) and glycerol (Fig. 2C) levels were also significantly lower in cachectic sera compared to uninfected controls, arguing against upregulation of lipolytic pathways. These data were consistent with our previous observation that the expression and phosphorylation of lipolytic enzymes hormone sensitive lipase (HSL), adipose triglyceride lipase (ATGL) and AKT were not increased in the vWAT, scWAT or liver of mice chronically infected with *T. gondii*^15^.

**Figure 2.**
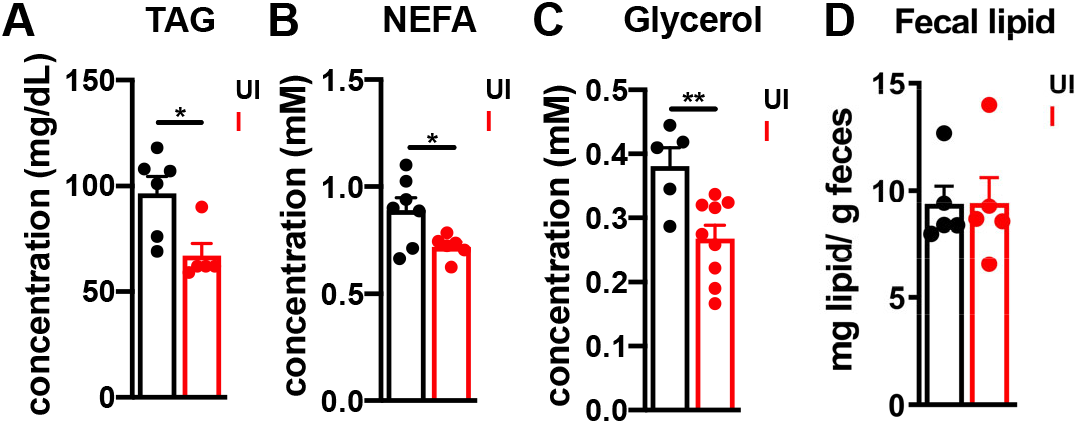
Chronically infected, cachectic mice have decreased lipids in blood but not in feces than uninfected littermates. At 9 weeks post-infection, mice were fasted for 4 hours and sera or fecal pellets were isolated. AC Serum triacylglyceride (TAG) (A), non-esterified fatty acids (NEFA) (B), glycerol (C) were measured by colorimetric assay. D, fecal lipids were precipitated and measured by dry weight. Data are normalized to uninfected. N = 5-7 mice per group from 2 independent experiments. Data represent the mean +/-SEM *p <0.05, **p<0.01 by unpaired Student t test.

The reduction in circulating triacylglycerides could be mediated by increasing storage, which primarily occurs through adipocyte hypertrophy and increased size. However, H&E staining indicated that adipocytes were significantly smaller in the vWAT of cachectic mice compared to uninfected controls (Figure S1A-B), consistent with a model wherein adipose depots rebound by hyperplasia (Figure 1C) rather than upregulating triglycerol storage. Circulating TAGs are primarily derived from the diet or synthesized in the liver from carbohydrates and free fatty acids^36^. The amount of lipid isolated from fecal pellets was similar between uninfected and infected mice, indicating that lipid absorption in the gut is not impaired in cachechtic mice (Fig. 2D). Cumulatively, these data are consistent with a model where the low serum TAG, NEFA and glycerol concentrations in cachetic mice are most likely the result of defective TAG synthesis in the liver, rather than impaired lipid uptake in the gut, increased storage in adipocytes or lipolysis.

### Serum glycerophospholipids and sphingolipids are dysregulated in *Toxoplasma* infection-induced cachectic mice

To understand whether the decrease in circulating lipids was generalizable across lipid species or specific to a subset of lipid families, sera was collected from cachectic, *T. gondii*-infected mice at 7 weeks post-infection or uninfected controls. The lipid metabolites in the sera were analyzed charged surface hybrid liquid chromatography-quadrupole time of flight mass spectrometry (CSH-QTOF MS). Of the 220 annotated lipids that were identified in the sera samples (Table S1), 33.18% were significantly differentially enriched (Fig. 3–4); the majority of which were underrepresented in infected, cachectic sera compaired to uninfected. Despite a decrease in total serum TAGs and NEFAs, cachectic mouse sera had a notable enrichment of some phosphatidylcholine (PC) and phosphatidylethanolamine (PE) species (Fig. 3, Fig. 4A), as well as increased ceramide and sphingomyelin species (Fig. 3, Fig. 4B), detected by positive and negative (Figure S2) ion modes. Consistent with the overall reduction of NEFA and glycerols (Fig. 2A-C), the annotated fatty acids (Fig. 4C), acylcarnitines (Fig. 4D), and triacylglycerols (Fig. 4E) were more abundant in uninfected sera relative to cachectic sera or not significantly different. The majority of steroids (Fig. 4F) were not significantly different between groups with the exception of cholesteryl easter (CE), 18:1 CE (oleate) and 22:6 CE were significantly enriched in the sera from infected, cachectic mice.

**Figure 3.**
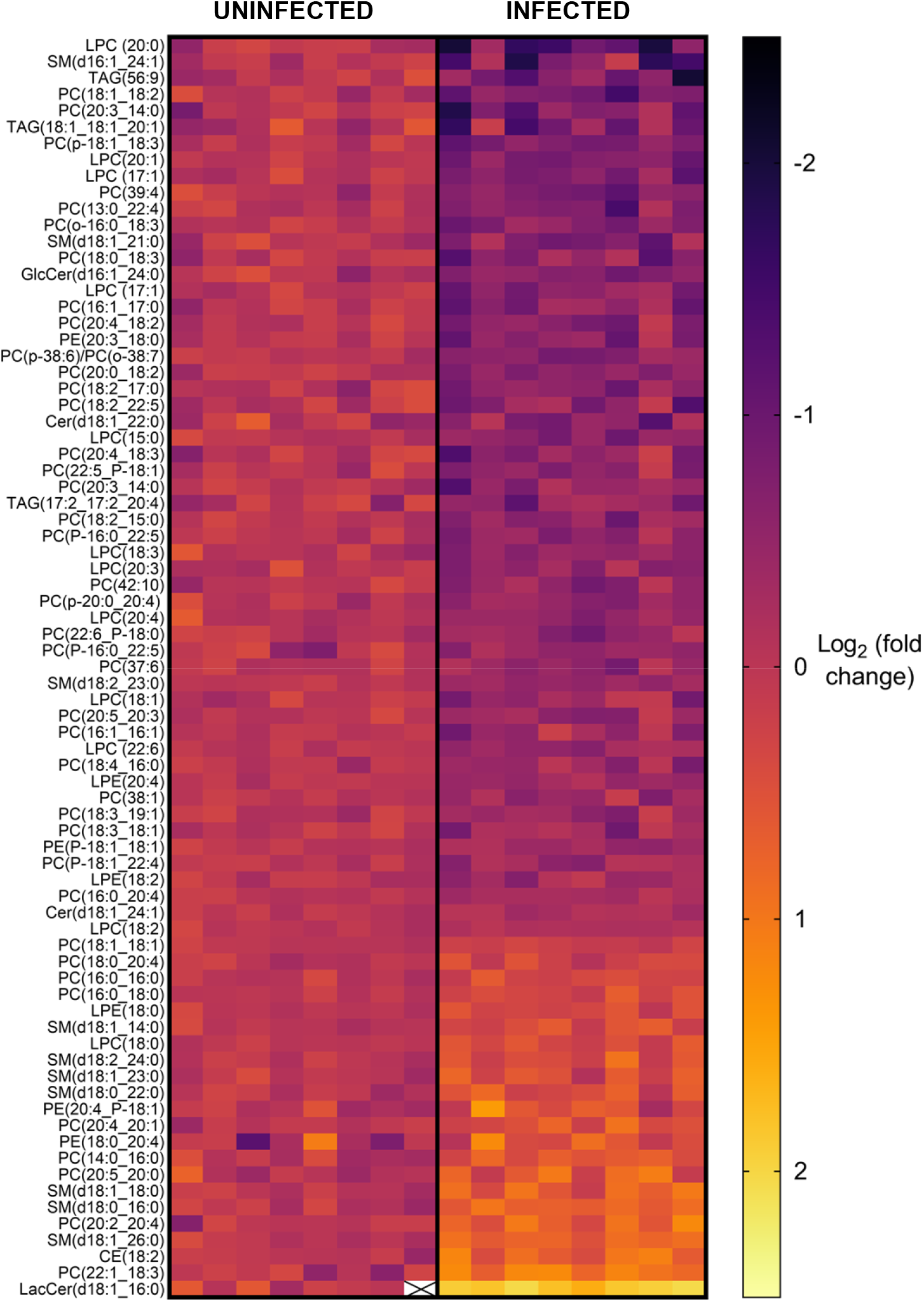
Chronically infected, cachectic mice have a distinct signature of circulating lipids relative to uninfected mice. 10-12 week old mice were infected intra-peritoneally with 10 Me49gLuc *T. gondii* cysts or mock injected with PBS. At 7 wpi sera was isolated retro-orbitally and untargeted analysis of complex lipids by CSH-QTOF MS/MS was performed. The heat map represents the 77 of 220 annotated lipids (Table S1) that were significantly differentially enriched by two-stage step-up method of Benjamini, Krieger, and Yekutieli with p<0.05. Each column is an individual mouse and the log2 (fold change) for each lipid specie (row) is displayed as the peak height observed in each mouse relative to the mean peak height of uninfected samples using ESI+ mode. LPC = lysophosphatidylcholine, Cer = ceramide, FA = fatty acids/fatty acyls, SM = sphingomyelin, PC = phosphatidylcholine, PE = phosphatidylethanolamine, GlcCer = glucosylceramide, LPE = lysophosphatidylethanolamine, TAG = triacylglyceride, CE = cholesteryl ester; LacCer = lactosylceramide. X represents missing values

**Figure 4.**
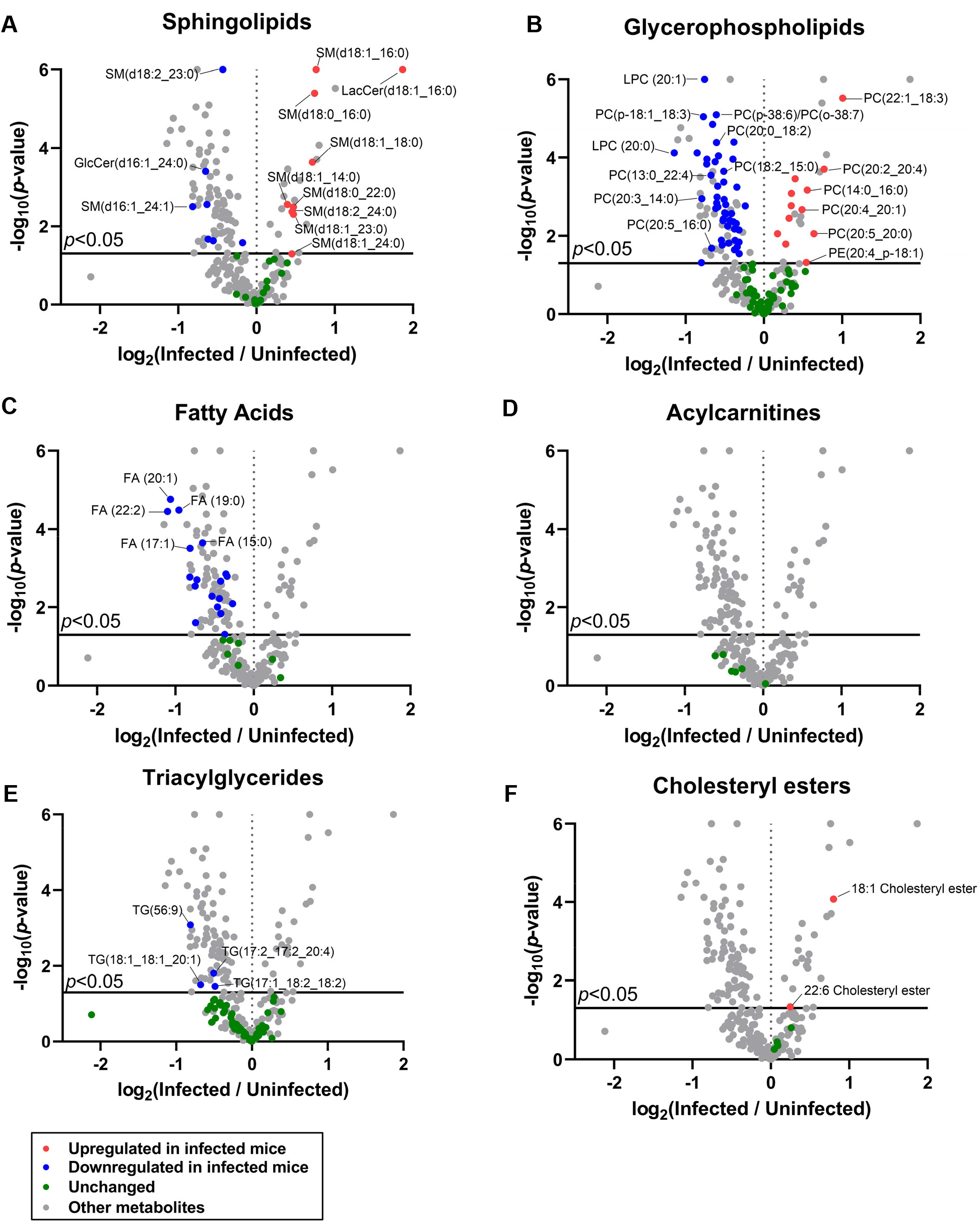
Chronically infected, cachectic mice have skewed lipid profiles in the blood. Each metabolites identified in Figure 4 represented as the mean peak height of infected sera relative uninfected samples, log2 (fold change). Y axis represents the −log_10_ of the *p*-value where *p* < 0.05 by unpaired Student’s t-test with False Discovery Rate of 1% by two-stage step-up method of Benjamini, Krieger, and Yekutieli. Metabolites are displayed by class: A, glycerophospholipids, B, fatty acids, C, acylcarnitine, D, triacylglyceride, E, cholesteryl esters. N = 8 mice per group.

Sphingolipids are major components of cell membranes with a range of signaling functions in immunity, metabolism, and tissue regeneration^37,38^. In cancer cachexia patients, a high circulating sphingolipid titer has recently been proposed as a marker of progressive cachexia pathology^32^. To more accurately measure changes in specific sphingolipid species we performed a targeted lipidomic analysis by LC-tandem quadrupole MS of sera from cachectic mice at 9 weeks post-infection relative to uninfected controls. Of the 34 sphingolipids measured, hexoceramide and long-chain bases (sphingoid bases) were enriched in the sera of cachectic mice relative to uninfected (Fig. 6A and Table S2), which was consistent with our total lipidomic analysis (Fig. 4–5).^39^. Long chain bases (LCBs) are the first products of de novo sphingolipid synthesis and deacylases, which can serve as precursors to complex sphingolipids^40^. All of the LCBs in our analysis were enriched in cachectic sera including sphingosine-1-phosphate (S1P), which has important signaling functions in cell survival, proliferation, differentiation and immune function; the S1P precursor sphingosine (Sph not significant), the most abundant mammalian long chain base; the saturated LCB, dihydrosphingosine (dhSph); and dihydrosphingosine-1-phosphate (dhS1P), which is generally represented as a minor LCB (Fig. 5B). Ceramides are N-acyl derivatives of long-chain bases and form the hydrophobic backbone of complex sphingolipids. C16-hexoceramide, C16-sphingomyelin, and C16-ceramide (not significant) were enriched in cachectic sera relative to uninfected (Fig. 5B). Notably, the very-long-chain ceramides and dihydroceramides were significantly depleted in cachectic sera relative to uninfected, consistent with emerging reports that phenotypes promoted by long-chain ceramides (C16 and C18) can be counter-regulated by very-long-chain ceramides (Fig. 6B, upper left quadrant)^41^.

**Figure 5.**
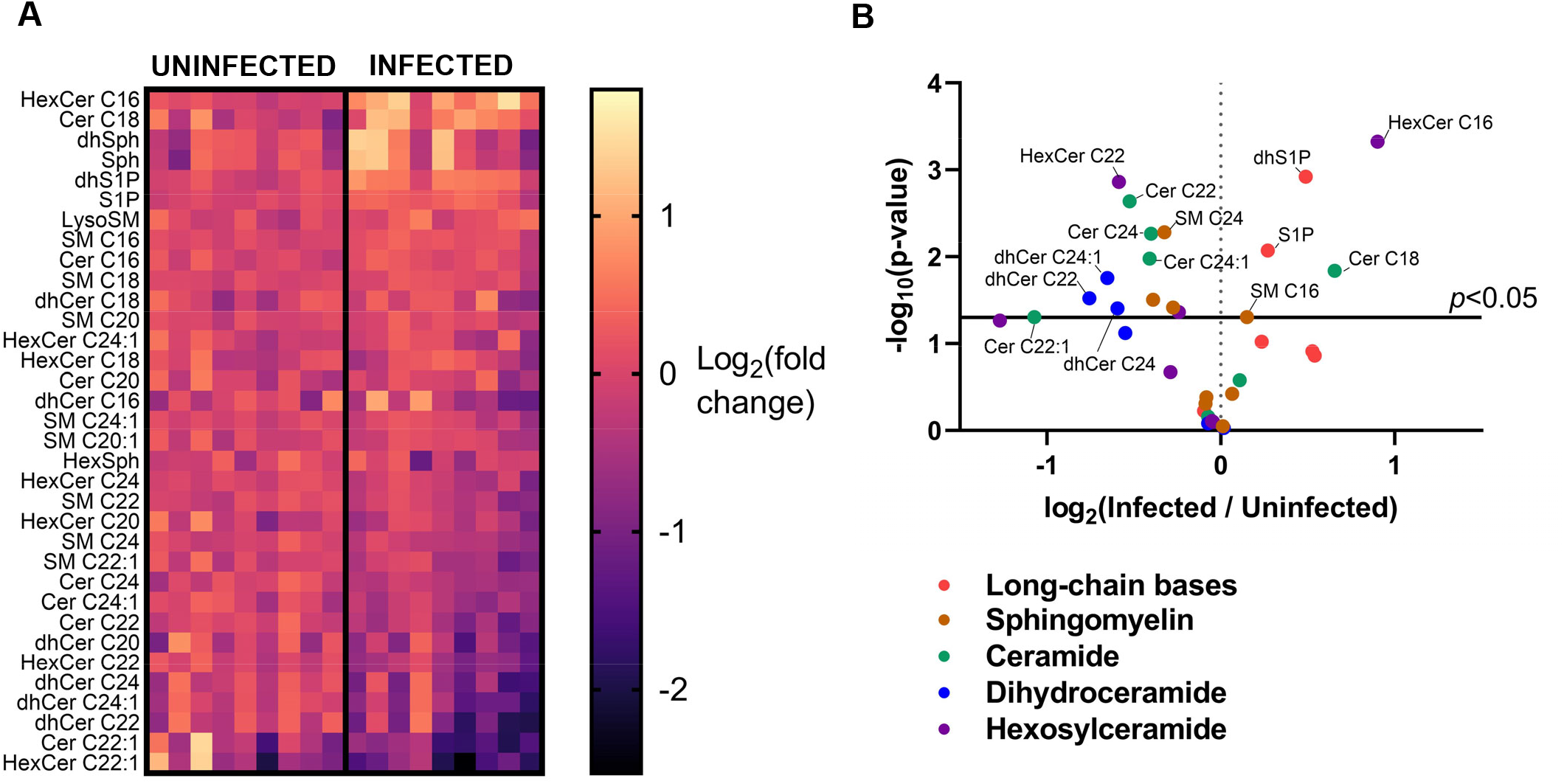
Significantly changed sphingolipids in the serum of cachectic mice compared to uninfected mice. Mice were intraperitoneally infected with 10 Me49gLuc *T. gondii* cysts of the strain or mock injected with PBS. At 9 wpi, mice sera was isolated retro-orbitally and sera analyzed by untargeted analysis of complex lipids by CSH-QTOF MS heat map in ESI + mode sphingolipids between infected and the corresponding control groups (n=9 mice per group). Normalized metabolite abundance (log2 transformed and row adjustment) are visualized as a color spectrum and the scale from least abundant to highest ranges is from −2.5 to 1.5. Black indicates low expression, whereas yellow indicates high expression of the detected metabolites. B, The volcano plot represents the mean peak height of infected sera relative uninfected samples, log_2_ (fold change). Y axis represents the -log_10_ of the *p*-value where *p* < 0.05 by unpaired Student’s t-test with False Discovery Rate of 1% by two-stage step-up method of Benjamini, Krieger, and Yekutieli. Cer=ceramide, HexCer= hexosylceramide, dhCer= dihydroceramide, SM=sphingomyelin.

### Cachectic mouse sera is deficient in TCA cycle intermediates downstream of α-ketogluterate

To evaluate a broader range of metabolic intermediates, the matching half of each sera sample from 7-weeks post infection (Fig. 3–4) was evaluated by automated liner exchange-cold injection system-gas chromatography-time-of-flight-mass spectrometry (ALEX-CIS GCMS-TOF). Of the 146 metabolites identified, 16.12% were significantly different between cachectic and uninfected sera (Fig. 7 and Table S3). Several liver metabolites associated were enriched in cachectic mouse sera including phenylalanine, a marker of liver dysfunction^42^. Increased serum and skeletal muscle phenylalanine has been observed in the C26 colon carcinoma model of cancer cachexia and was associated with decreased food intake^43,44^. Fructose, which is metabolized in the liver, was also elevated in the sera of cachectic mice. Fructose has been shown to regulate the muscle specific E3 ubiquitin ligases Murf-1 and Atrogin-1 which drive muscle atrophy during cachexia^45^. We also observed increased abundance of diet-derived metabolites including the antiinflammatory di-saccharide trehalose; pinatol, a plant methylated inositol; and the bacterial metabolite L-ribonate, which has been observed in the sera of diabetic mice after exercise^46^.

Confirming our measurement of total serum glycerol at 9 wpi (Fig. 3), glycerol was significantly reduced in the metabolomic analysis of cachectic mouse sera (Fig. 6A). Consistent with our previous report that infected, cachectic mice have low glucose levels but remain responsive to insulin^15^, glucose was slightly but significantly reduced in cachectic sera, however, its immediate glycolytic product glucose-6-phosphate (G6P) was comparable in between experimental groups (Fig. 6B).

**Figure 6.**
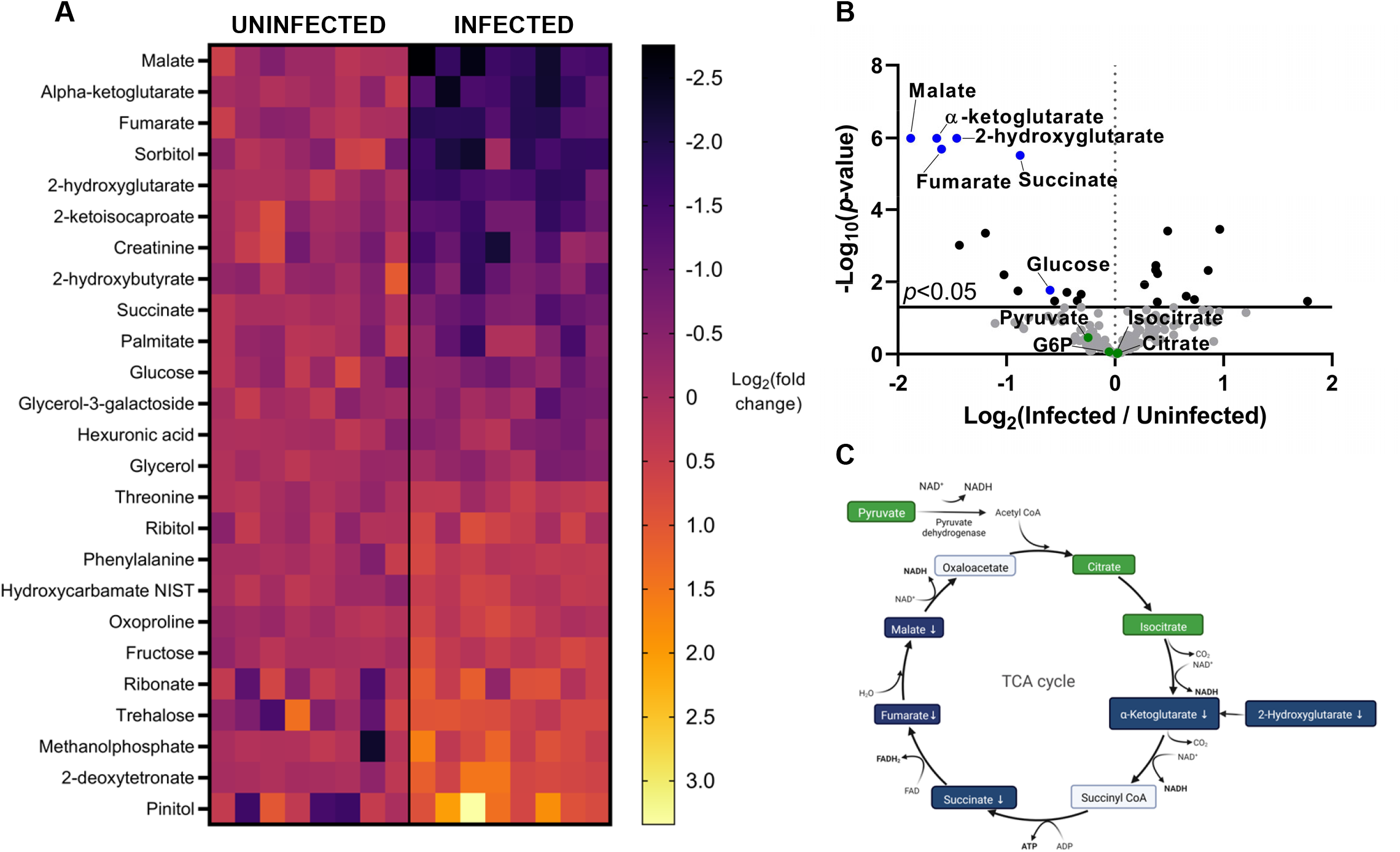
Cachectic mice have are deficient in circulating TCA cycle intermediates. Untargeted serum metabolites analysis (ALEX-CIS GCTOF-MS) was performed on the same samples analyzed in Figure 4. **A**, Heat map represents significantly differentially enriched metabolites where each column is a sample from an individual mouse. The Log2 (fold change) for each metabolite (row) is displayed as the peak height observed in each mouse relative to the mean peak height of uninfected samples using ESI+ mode. Ranked from least abundant in Infected animal (−2.5, Black) to most abundant (2.5, yellow). **B**, Volcano plots represent log2 (fold change) of the mean peak height for infected mice relative to uninfected mice. TCA cycle intermediates are annotated. *P* < 0.05 by unpaired Student’s t-test with False Discovery Rate of 1% by two-stage step-up method of Benjamini, Krieger, and Yekutieli. **C**, Schematic of metabolic dysregulation of the TCA cycle, where green represents metabolites that were detected but not significantly differentially enriched between uninfected and infected, cachectic mice, grey represents metabolites that were not detected, and blue represents metabolites that were significantly less abundant in infected, cachectic sera.

The TCA cycle intermediates malate, α-ketoglutarate, fumarate, 2-hydroxyglutarate, and succinate represented a cluster of the most highly depleted metabolites in cachectic sera relative to uninfected sera (Fig. 6A-B). Other TCA cycle intermediates pyruvate, citrate, and isocitrate were detected, however, the levels were not significantly different between uninfected and cachectic sera (Fig. 6A-B). Together these data suggest a blockade in the TCA cycle at isocitrate dehydrogenase which converts isocitrate to α-ketoglutarate in chronic *T. gondii* infection-induced cachexia (Fig. 6C).

## Discussion

The liver is a central regulator of lipid homeostasis in mammals. The reduction of circulating NEFA, glycerol, and TAGs in mice with *T. gondii*-induced chronic cachexia led to the hypothesis that the dysregulated lipid metabolism is involved in the pathology of chronic cachexia. Our data are consistent with a model where liver atrophy-associated dysregulated lipid metabolism aggravates the imbalanced energy metabolism in mice with *T. gondii*-induced chronic cachexia. The liver plays indispensable roles in nutrient metabolism, lipid homeostasis, and the immune response, dysregulation of these processes during hepatic disorders causes systemic wasting^47^. Patients with chronic liver disease have reduced muscle mass^48^. Kurosawa *et al*. used the bile duct ligation model of liver injury to demonstrate a causal relationship between liver fibrosis and muscle wasting^49^, highlighting the importance of liver function in systemic metabolism. Defective lipid metabolism in the liver is becoming accepted as a contributor to peripheral tissue wasting in cancer cachexia^18^, underlining the importance of lipid metabolism in the pathology of cachexia.

Consistent with the reduction in total serum NEFAs, glycerol and TAGs, we observed less diversity in lipid species in cachectic, infected mouse sera compared to uninfected controls. Sphingolipids, which were among the minority of species enriched in cachectic sera, are increasingly recognized as disease biomarkers in a range of metabolic, cardiovascular, inflammatory, and neurological disorders^50,51^. Using LC–MS, Morigny *et al*. determined that levels of C16 sphingomyelin (SM), C16 ceramide (CER), and C16 hexosylceramide (HCER) in the plasma were positively correlated with weight loss in gastric cancer patients^32^. Similar to these results, C16 SMs were enriched in *T. gondii-infected*, cachectic mice at 9 weeks post-infection compared to non-infected mice (Figure 5). These data indicate that sphingolipid dysregulation may be a common driver of cachexia across disease etiologies and underscore the clinical relevance of *T. gondii*-induced cachexia model.

In clinical studies, sphingolipid biosynthesis has been shown to be positively regulated by tumor necrosis factor-alpha (TNF-α) by this cytokine’s ability to upregulate sphingomyelinase (SMase) activity^52,53^. Consistent with this potential mechanism, our lab and others have shown that mice with chronic *T. gondii* infection sustain low but significantly elevated levels of sera TNF-α^15^. It is also possible that this dysregulated sphingolipid metabolism promotes inflammatory signaling in *T. gondii-induced* cachexia. For example, long-chain C16-C18 ceramides have been shown to induce nuclear translocation of NF-Kb^54,55^, which may explain our observation that NF-Kb-dependent transcripts TNF-α, IL-12, IL-1, and IL-6 are elevated in circulation during *T. gondii-* induced cachexia long after systemic parasite infection is cleared^15^. Lactosylceramide (LacCer), another significantly sphingolipid in the lipidomic study, is able to be induced by TNF-α stimulation. LacCer has been shown as an inducer of oxidative stress and prostaglandins precursor arachidonic acid^39^, which may propagate inflammatory signaling in the cachectic mice.

In a sphingolipid-targeted analysis, we determined that the LCBs (Figure 6) were uniformly elevated in cachectic mouse sera relative to uninfected mice. LCBs are synthesized de novo in the ER from serine and acyl CoA, diacylation of ceramides, or derived from circulating S1P and dietary sphingolipids processing in the endolysosome^40^. We have previously shown that the commensal microbiota of mice with *T. gondii*-induced cachexia is disrupted during chronic infection^26^. Dysbiotic commensal microbiota could be a source of LCBs we observed in the sera of infected, cachectic animals, which would be consistent with the differential enrichment of commensal and diet metabolites relative to uninfected mouse sera (Figure 5). While it is formally possible that the increased LCB species are due to changes in intestinal absorption during cachexia, our data indicating that lipids were shed at equivalent levels in the feces of uninfected and infected cachectic mice (Figure 2D) argue against this, as does previously published data showing an equivalent number of calories in the feces^15^. SPH (or dhSPH) is most frequently formed by the loss of the acyl group from ceramide (or dhCer) by a gain of ceramidase activity or loss of desaturase activity. So our overall reduction on ceramides is consistent with a models where de novo synthesis of LCBs is increased (Figure 2), although metabolic flux experiments would be necessary to determine this definitively.

Interestingly, our data showed a striking reduction in TCA cycle intermediates alphaketogluterate, malate, fumerate, succinate, and 2-hydroxyglutarate. Consistent with a significant shift in mitochondrial metabolism, sphingolipids are critical structural components of the mitochondrial membrane, regulating mitochondrial homeostasis by interacting with integral membrane proteins and mitochondrial regulatory machinery^30,56^. For example, C18 ceramide facilitates the anchoring of LC3B-II autophagosomes on outer mitochondrial membrane thereby promoting mitophagy^57,58^. In eukaryotic cells, the de novo biosynthesis pathway is a major source of mitochondrial sphingolipids^31^. In addition, mitochondria are capable to synthesize sphingolipids using ceramide synthase^31^. Several lines of evidence suggest that dysregulation of sphingolipid metabolism is detrimental to mitochondrial function. Sphingolipids accumulation in mitochondria is associated with increased mitochondrial reactive oxygen species (ROS) and reduced ATP production in patients with metabolic disorders^31^. Similarly, Knupp *et al*. demonstrate a mechanism where increased sphingolipid synthesis in yeasts lead to dysregulated electron transport chain (ETC) activity in mitochondria, ROS accumulation, and cell death in a nutrient-dependent manner^30^.

Intriguingly, there is growing recognition of the association between dysregulated sphingolipid metabolism and mitochondrial disorders in liver dysfunction, including liver fibrosis, insulin-resistant liver, and non-alcoholic fatty liver disease^31,59^. Although this relationship has not been explored in cachexia directly, cachectic mice have been reported to have decreased levels of mitochondrial fusion markers (mitofusin-1, mitofusin-2, and opa-1), reduced biogenesis marker (peroxisome proliferator-activated receptor-gamma coactivator), and increased mitophagy marker (p62)^60^ consistent with disrupted mitochondrial homeostasis. These lines of evidence combined with our data showing liver atrophy and fibrosis, serum sphingolipid enrichment, and reduction in TCA cycle intermediates suggest a model where mice dysregulated sphingolipid metabolism drives mitochondrial metabolic dysfunction in the liver. Metabolic flux experiments will be necessary to determine the source of elevated sphingolipids, and whether there is a causal link to mitochondrial dysfunction in the liver or other tissues.

The second subset of lipid species enriched in cachectic sera was a subset of phosphatidylcholines (PCs). PCs are the critical component of mammalian cell membranes^61^, signaling molecule precursors and the major phospholipid in bile^61^. Glucose metabolism influences PC synthesis the nucleotide end products of the pentose phosphate pathway (PPP) and diacylglycerol (DAG) are necessary to produce cytidine diphosphate choline and convert it into PC, respectively^62^. Consistent with our previous report^15^, glucose was significantly decreased in cachectic sera (Fig.7A-B). Reduced glucose could limit flux through the PPP necessary to generate precursors for PC synthesis in *T. gondii*-induced chronic cachexia. It is also possible that reduced flux through the TCA cycle could promote sphingolipid synthesis. Serine, the building block for sphingolipids, is synthesized from a glycolytic intermediate 3-phosphoglycerate (3PG)^62^. Blockade of the TCA cycle lead to substrate availability for serine synthesis and the subsequent *de novo* sphingolipid biosynthesis in mice with *T. gondii*-induced cachexia. However, the source of PC substrates remains to be determined by metabolic flux experiments. It is notable that in skeletal muscle, PC catabolism is necessary for mitochondrial biogenesis and oxidative capacity, although this relationship has been more extensively studied in non-cachexia scenario^63,64^.

Whether the cachexia associate metabolic dysregulation benefits *T. gondii*, mediates host restriction of the parasite or primarily provokes pathology is an open question. The majority of previous metabolomic studies in *T. gondii* infection were performed during the acute or acute to chronic phase transition of infection and cachexia was not included in the criteria for assessment. Our data demonstrate that *T. gondii*-induced chronic cachexia shares metabolic signatures in cachectic patients. By pairing metabolomic and lipidomic analyses, our data point to areas of crosstalk between PC biogenesis, sphingolipid metabolism and dysregulated TCA cycle signaling that may synergize to provoke metabolic dysfunction during *T. gondii* infection-induced cachexia. This study is part of a growing body of literature supporting a role for dysregulated lipid metabolism in the pathogenesis of cachexia conserved across an infection, liver disease, and cancer-induced etiologies of the disease, indicating that this may be an important area for therapeutic development.

## Limitation of the study

This study relied on metabolic analysis of sera at discrete time points in infection-induced cachexia. Future studies using labeled substrates will be necessary to determine which metabolic pathways are primarily disrupted in this model. Pairing these experiments with measurements in discrete tissues including the muscle, liver, and adipose tissue will be necessary to determine which tissue sites are primarily driving the mechanism of metabolic disruption in T. gondii-induced cachexia.

## Author contributions

TYF: lipidomic, metabolomics and histological data analysis, manuscript preparation; SJM: study conceptualization, *in vivo* experimentation, data analysis and manuscript preparation; HG: lipidomic and metabolimics data analysis; TEF: targeted sphingolipid experimentation and analysis; MK: lipidomics experiment conceptualization.

## Acknowledgments

The authors wish to acknowleadge the technical support from the National Institute of Health (NIH) West Coast Metabolomics Center at the University of California, Davis. In addition, we would like to thank Dr. Bochen Yin for advice on volcano plot analysis and Dr. Farzana Begum Liakath for the preliminary assessment of sphingolipid data.

## STAR Methods

### Mice

CBA/J (RRID:IMSR_JAX:000656) and C57BL/6 (RRID:IMSR_JAX:000664) mice were purchased from Jackson Laboratories. Mice were maintained, infected and monitored in accordance with the University of Virginia Institutional Animal Care and Use Committee AAALAC and IACUC protocol #4107-12-18.

### Infections

To generate cysts for infection, 8-10 week female CBA/J mice were infected with 10 Me49 bradyzoite cysts by intraperitoneal injection. 4–8 weeks following infection, mice were euthanized by CO2 inhalation and cysts were harvest from brains homogenate passed through a 70 μm filter. Homegenate was washed 3 times in PBS, stained with dolichos biflorus agglutinin conjugated to FITC (Vector labs) at a 1:500 dilution. The number of cysts were determined by counting FITC-positive cysts at 20x magnification using an EVOS FL imaging system (Thermo Fisher). For experimental infections 10–14-week-old male C57BL/6 mice were infected with 10 Me49 bradyzoite cysts by intraperitoneal infection resuspended in 200 Ml PBS per mouse using a 5G 5/8" tuberculin syringe. Prior to infection, mice were cross-housed on dirty, wood chip bedding for two weeks to normalize commensal microbiota and limit the effect of eating corn husk bedding on dietary metabolites. At experimental endpoints, mice were fasted for 4 hours and isoflurane anaesthetized to isolate sera via retro-orbital bleed and/or euthanized by CO_2_ asphyxiation to harvest tissues for weighing and histological analysis.

Chronic parasite burden was determined by Qpcr of *T. gondii* 529bp Repeat Element (RE) (*forward:* 5’-CACAGAAGGGACAGAAGTCGAA-3’; *reverse:* 5’-CAGTCCTGATATCTCTCCTCCAAGA-3’; *probe:* 5’-CTACAGACGCGATGCC-3’) compared to mouse beta actin Mm02619580_g (ThermoFisher Scientific) as described^65^ and analyzed using the ΔCt method. Brain DNA was isolated as described^66^, and used at 100 ng DNA per Qpcr reaction on a QuantStudio6 instrument (Applied Biosystems).

### Tissue weights and histological examination

At 9 weeks post infection mice were euthanized with CO2 asphyxiation and blood was isolated by cardiac exsanguination. Abdominal subcutaneous white adipose depots, epididymal visceral white adipose depots, and, quadricep muscles were isolated and weighed. Alternatively, adipose tissue was formalin fixed for 24 hours, transferred to sucrose and embedded in optimal cutting temperature compound (OTC) for cryosectioning. Sections were stained H&E at the University of Virginia Research Histology core. The stained fat tissue sections were imaged with a Slide Scanner (Leica) and images were processed using the QuPath 0.3.2^67^. Adipocytes on the slides were detected by the in-built classifier in the QuPath according to the method developed by Palomaki *et al*.^68^. Briefly, a random trees-based pixel classifier was trained to detect the adipocytes cells by the morphology. To optimized the adipocyte detection, all the fat tissue sections in this study were used in the training. The final pixel classifier was applied to all the fat slides to detect the adipocytes across the whole section with the following setting: min object size 20 μm^2^, min hole size 30 μm^2^. The area for each detected adipocyte was measured by in-built tools in the QuPath.

### Sera metabolomics and lipidomics

At 7 weeks post-infection, isoflurane-anesthetized mice were retro-orbitally bled and sera was flash frozen and sent to the National Institute of Health (NIH) West Coast Metabolomics Center (UC Davis) for untargeted mass spectrometry analysisusing the primary metabolism assay (ALEX-CIS GCTOF-MS) or the complex lipids (CSH-QTOF MS) assay. Detected meatbolites were identified based on retention time and mass spectra from MassBank of North America, curated by the NIH West Coast Metabolomics Center, and reported as raw peak heights (Table S1 and S3). The raw peak heights from each analytical platform were normalized to the average peak heights of the identified metabolites in uninfected group. The resulting data were analyzed for fold-change and multiple unpaired t-test and visualized using volcano plots to identify the differential expression of metabolites in response to *T. gondii-induced* cachexia.

Lipid extraction and analysis for targeted sphingolipid analysis was done using liquid chromatography-electrospray ionization-tandem mass spectrometry (LC-ESI-MS/MS)^69^. Lipids were extracted from sera using an azeotrophic mix of isopropanol:water:ethyl acetate (3:1:6; v:v:v). Internal standards (10 pmol of d17 long-chain bases and C12 acylated sphingolipids) were added to samples at the onset of the extraction procedure. Extracts were separated on a Waters I-class Acquity UPLC chromatography system. Mobile phases were (A) 60:40 water:acetonitrile and (B) 90:10 isopropanol:methanol with both mobile phases containing 5 mM ammonium formate and 0.1% formic acid. A Waters C18 CSH 2.1 mm ID × 10 cm column maintained at 65°C was used for the separation of the sphingoid bases, 1-phosphates, and acylated sphingolipids. The eluate was analyzed with an inline Waters TQ-S mass spectrometer using multiple reaction monitoring.

### Statistical analysis

Statistical analysis was performed using GraphPad Prism 9 software (GraphPad). Statisticall differences between infected, cachectic mice and uninfected mice were determined using unpaired *t*-test, as indicated in the figure legend. *P* value <0.05 was considered significant.

