## Supplemental figures and legend for "Tricarboxylic acid (TCA) cycle, sphingolipid, and phosphatidylcholine metabolism are dysregulated in *T. gondii* infection-induced cachexia"

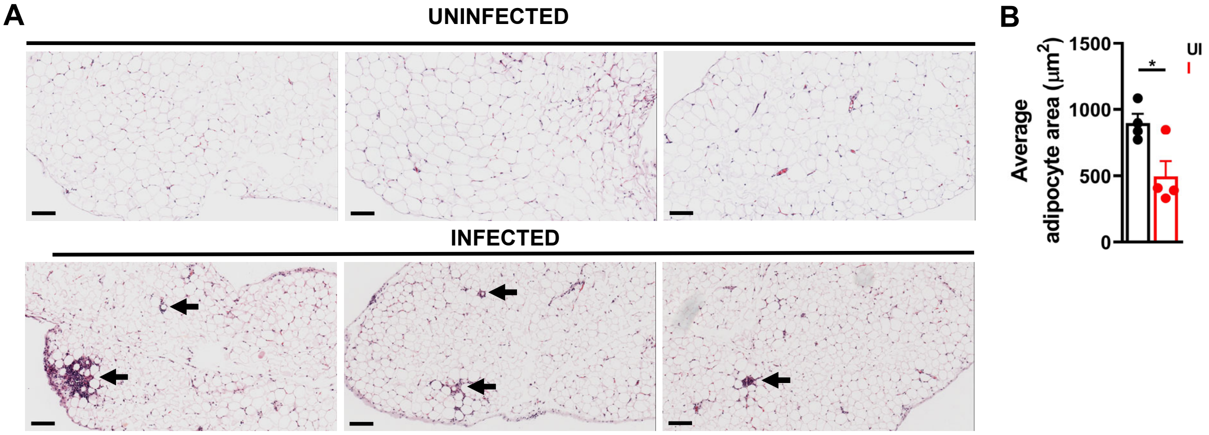


**Figure S1. Mice with *T. gondii*-induced cachexia have smaller adipocytes compared to uninfected, non-cachectic mice. A,** At nine weeks post-infection, formalin-fixed paraffin-embedded adipose tissues were stained with hematoxylin and eosin. Arrows indicate crown-like structures. Scale bar= 100 mm **B**, Average adipocyte area across a whole section of vWAT was measured using QuPath-0.3.2. Each symbol represents the average adipocyte area from an individual mouse. N=4 mice per group from 2 independent experiments. Error bars represent the mean +/- SEM. *$p < 0.05$ by unpaired Student's t-test.

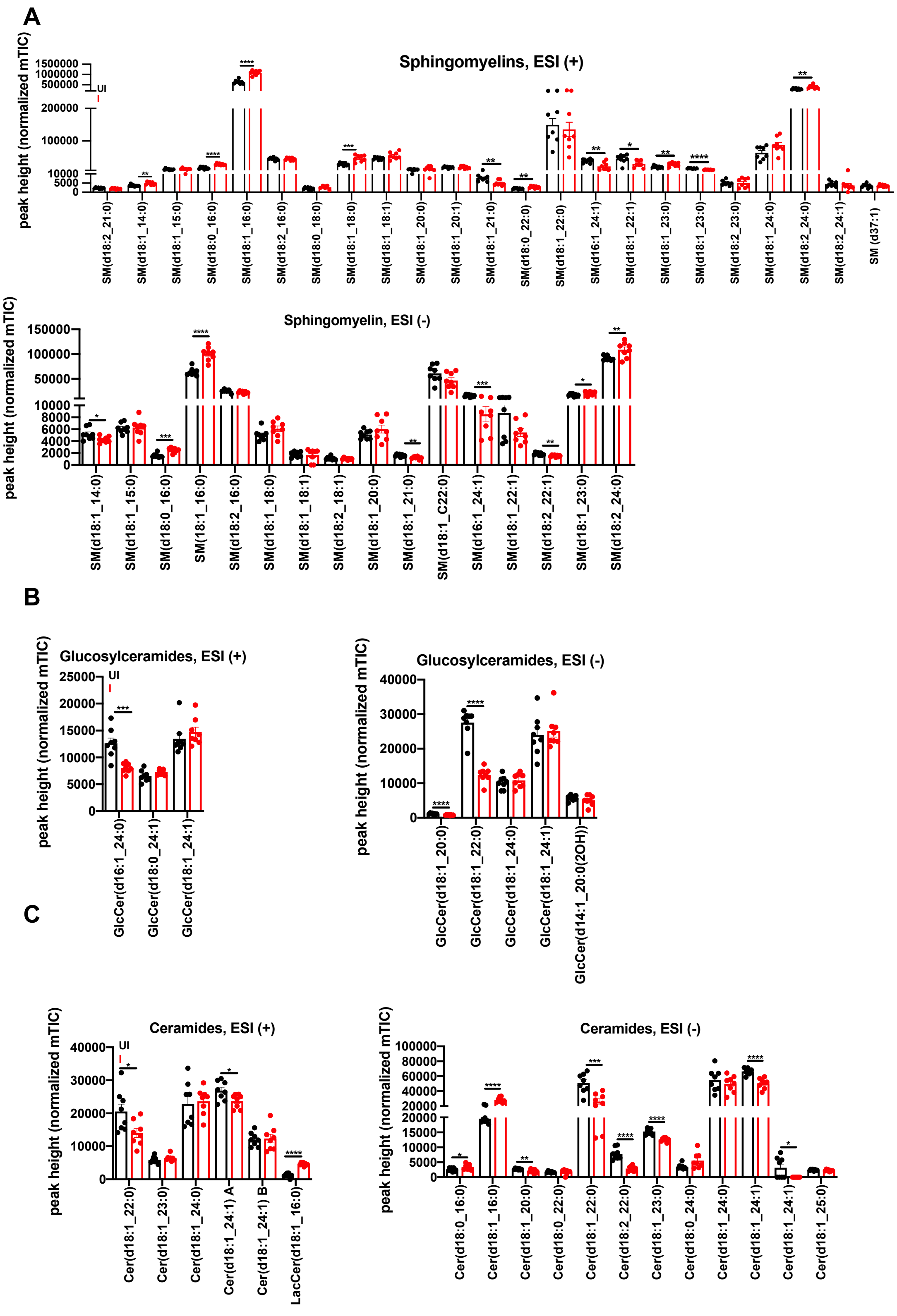


**Figure S2. The detected changes of sphingolipids from cachectic mice in ESI positive mode are similar to those in ESI negative mode .** Mice were intraperitoneally infected with 10 Me49gLuc *T. gondii* cysts of the strain or mock injected with PBS. At 9 weeks post infection, mice sera (n = 8 mice per group ) were isolated retro-orbitally and sera analyzed by untargeted analysis of complex lipids by CSH-QTOF MS/MS **A-C,** Data is represented as ‘norm mTIC’ which is relative semi-quantifications of sphingolipids in positive and negative ion mode for sphingomyelins (**A**), glucosylceramides (**B**), and ceramides (**C**). Each symbol represents an individual mouse. Data represent the mean +/- SEM. *$p$<0.05, **$p$<0.01, ***$p$<0.001 by unpaired Student’s t-test.

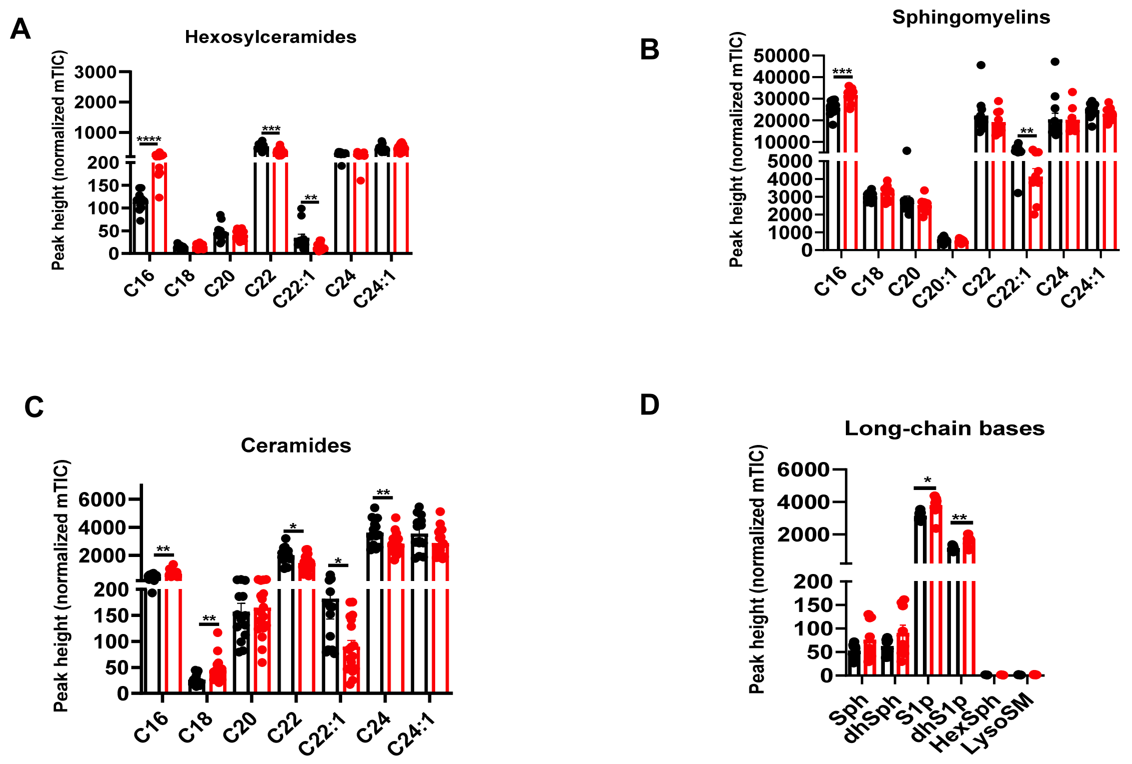


**Figure S3. Cachectic mice with chronic *T. gondii* infection have significant dysregulation in circulating sphingolipids species compared to uninfected mice.** 10-14 week old C57BL/6J mice were intraperitoneally infected with 10 Me49gLuc *T. gondii* cysts of the strain or mock injected with PBS. At 9 weeks post infection, mice sera were isolated retro-orbitally and sera analyzed by untargeted analysis of complex lipids by CSH-QTOF MS/MS A-D, each sphingolipid class is represented as relative semi-quantifications normalized mTIC of each class. (**A**), Hexocylceramide (**B)** sphingomyelin**,** (**C**) Ceramides **(D)** Long-chain bases. Each point represents a mouse. n =9 mice per group. *Each point represents an idividual mouse. Data represent the mean +/- SEM. *p<0.05, ***p*<0.01, ****p*<0.001 by unpaired Student's t-test.
